# Translational activation by an alternative sigma factor in *Bacillus subtilis*

**DOI:** 10.1101/2021.03.06.434181

**Authors:** Dylan M. McCormick, Jean-Benoît Lalanne, Tammy C. T. Lan, Silvi Rouskin, Gene-Wei Li

## Abstract

Sigma factors are an important class of bacterial transcription factors that lend specificity to RNA polymerases by binding to distinct promoter elements for genes in their regulons. Here we show that activation of the general stress sigma factor, σ^B^, in *Bacillus subtilis* paradoxically leads to dramatic induction of translation for a subset of its regulon genes. These genes are translationally repressed when transcribed by the housekeeping sigma factor, σ^A^, owing to extended RNA secondary structures as determined *in vivo* using DMS-MaPseq. Transcription from σ^B^-dependent promoters liberates the secondary structures and activates translation, leading to dual induction. Translation efficiencies between σ^B^- and σ^A^-dependent RNA isoforms can vary by up to 100-fold, which in multiple cases exceeds the magnitude of transcriptional induction. These results highlight the role of long-range RNA folding in modulating translation and demonstrate that a transcription factor can regulate protein synthesis beyond its effects on transcript levels.

## INTRODUCTION

Transcriptional regulation by sigma factors is a hallmark of bacterial gene expression. Sigma factors bind to the core RNA polymerases, forming holoenzymes that can initiate transcription at sites with well-defined sequences. In *B. subtilis*, most genes are transcribed by the housekeeping sigma factor σ^A^, and some are additionally or exclusively transcribed by alternative sigma factors that control specific processes such as sporulation and motility (Haldenwang 1995; Helmann 2019). The alternative sigma factor σ^B^ is involved in the general stress response (Haldenwang and Losick 1979; Hecker et al. 2007; Price 2014; Haldenwang 1995) and initiates transcription for over two hundred genes with well-defined promoter sequences (Nicolas et al. 2012; Petersohn et al. 1999; Zhu and Stülke 2018). Induction of transcription leads to corresponding increases in RNA levels (Figure 1A).

**Figure 1.**
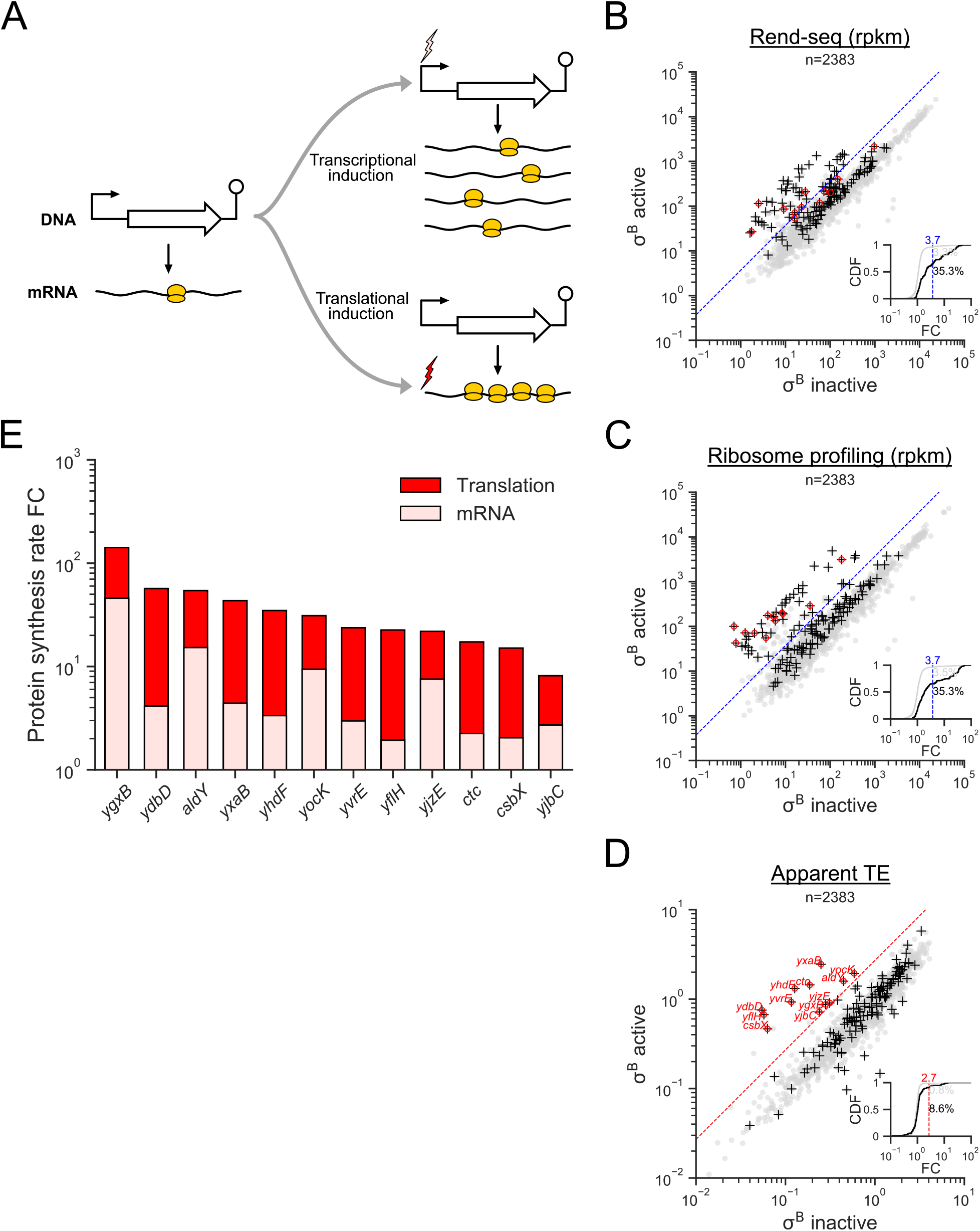
σ^B^ activates both transcription and translation. (**A**) Models of transcriptional and translational induction for a transcriptional unit consisting of a promoter, coding sequence, and terminator. Stimuli are indicated with lightning bolts and ribosomes are colored in yellow. (**B**) RNA-seq, (**C**) ribosome profiling, and (**D**) apparent translation efficiency measurements from σ^B^ active and inactive conditions. σ^B^ regulon genes are indicated with black crosses (+), and a subset that are translationally activated are highlighted in red. The dashed blue lines mark a 3.7-fold change in expression for visual reference. The dashed red line is an approximate threshold (2.7-fold) separating the population of translationally activated genes from those whose apparent TE does not markedly change. The insets show the cumulative distribution function (CDF) of fold change (FC) across the two conditions in each measurement, with separate CDFs for all genes (gray) and σ^B^ regulon genes (black). The percentage of genes in each group exceeding the chosen thresholds are listed on the right. (**E**) Contributions of mRNA levels and translation to changes in protein synthesis rate among translationally activated σ^B^ regulon genes. The fold change in protein synthesis rate is indicated by the height of the bars, with the light and dark red regions denoting the respective contributions of mRNA levels and translation, i.e., fold-change in protein synthesis = (fold-change in mRNA level)×(fold-change in translation efficiency).

Translational regulation is also widespread in *B. subtilis*, although it is not typically thought to be controlled by transcription factors. Differential translation among genes in the same operon is largely driven by differences in mRNA secondary structure (Burkhardt et al. 2017) and is important for stoichiometric production of proteins in the same complex or metabolic pathway (Lalanne et al. 2018; Li et al. 2014). Translation can be additionally regulated by RNA-binding proteins or riboswitches that modulate the accessibility of the ribosome binding sites on the mRNA (Breaker 2018; Yakhnin et al. 2004, 2007). Operons are often controlled both transcriptionally and translationally (Figure 1A), but seldomly by the same regulator (Bastet et al. 2018; Chauvier et al. 2017; Hollands et al. 2012).

Here we show that the transcription factor σ^B^ not only activates transcription, but also derepresses translation for a subset of its regulon genes. Using Rend-seq (end-enriched RNA-seq) (Lalanne et al. 2018) and ribosome profiling, we identified 12 genes whose apparent translation efficiency is increased substantially during σ^B^ activation. Most of them are transcribed from a σ^B^-dependent promoter as well as at least one σ^A^-dependent promoter, generating multiple transcript isoforms. By modulating σ^B^ activities, we found that each transcript isoform is associated with a distinct translation efficiency, with strongly repressed translation for σ^A^-driven isoforms and elevated translation for σ^B^-driven isoforms. These were orthogonally confirmed using a fluorescent reporter in a subset of examples. Both computational RNA folding and *in vivo* structural probing by DMS-MaPseq (Zubradt et al. 2016) indicate that the repressed σ^A^-driven isoforms possess extended RNA secondary structures that sequester the ribosome binding sites. On the other hand, σ^B^-driven isoforms have shorter 5’ UTRs that only include the regions corresponding to the second halves of the extended stem-loops in the longer σ^A^-driven isoforms. Therefore, σ^B^ can simultaneously activate both transcription and translation by modulating isoform-specific secondary structures.

## RESULTS

### σ^B^ activates translation for a subset of its regulon

We first observed translational activation of σ^B^ regulon genes while profiling gene expression for a *B. subtilis* strain with an elevated general stress response during steady-state growth due to a genetic modification (Methods). Rend-seq and ribosome profiling data were generated to quantify the mRNA levels and protein synthesis rates, respectively, for both the wild type (“σ^B^ inactive”) and the genetically modified strain (“σ^B^ active”). The density of ribosome footprints of a gene provides an estimate for the relative rate of protein synthesis, provided that most ribosomes complete translation to yield full-length polypeptides and that the elongation time averaged across the entire transcript is constant (Ingolia et al. 2009; Li 2015; Lalanne et al. 2018; Li et al. 2014). Translation efficiency (TE), defined as the rate of protein production per mRNA molecule, can then be estimated from Rend-seq and ribosome profiling data by calculating the per-gene ribosome profiling coverage over Rend-seq coverage, i.e., the ribosome density along a transcript (Li 2015; Li et al. 2014). Given σ^B^’s well-understood role in transcription initiation, we expected its regulon members to change in mRNA levels and not TE.

Surprisingly, we found that several genes in the σ^B^ regulon showed far greater increases in protein synthesis rate (ribosome profiling) than in mRNA levels (Rend-seq). Between the two conditions, 25% of the annotated σ^B^ regulon genes (Zhu and Stülke 2018) had substantially different expression levels (56/225 with >3.7-fold change, Figure 1B and 1C). Although most genes showed concordant changes in mRNA levels and protein synthesis rates, a notable population (21%, 12/56) exhibited a considerably greater increase in protein synthesis rates than mRNA levels (>2.7-fold), suggesting an increase in apparent translation efficiency (Figure 1D). Among these translationally activated σ^B^ regulon genes, the magnitude of TE increases often exceeded the rise in mRNA levels, as most genes (75%, 9/12) exhibited a fold change in apparent TE accounting for >50% of the observed fold change in protein synthesis rate (Figure 1E). Hence, translational induction contributes to the majority of the increase in expression of a subset of the σ^B^ regulon, suggesting a yet-unknown strategy for activating translation following σ^B^ induction.

### σ^B^-dependent alternative mRNA isoforms drive translational upregulation

To identify the regulatory features that could drive translational upregulation, we examined the transcript architecture of translationally activated σ^B^ regulon genes using Rend-seq. Through sparse fragmentation of input RNAs, Rend-seq enriches for the 5’ and 3’ boundaries of transcripts, enabling the detection and quantification of mRNA isoforms within operons (Lalanne et al. 2018). We observed that the translationally activated σ^B^ regulon genes were found in two or more different RNA isoforms (Figure 2, Figure S1, Figure S2). In particular, 8 of the 12 genes shared a common operon architecture (Figure 2, Figure S1): They were each transcribed both as a part of a polycistronic mRNA from a vegetative (σ^A^-dependent) promoter, as well as from their own σ^B^-dependent promoter. As illustrated by the representative genes *ctc* and *yvrE*, in the absence of stress, the primary isoform was the long, σ^A^-dependent polycistronic mRNA (Figure 2). In these transcripts, the ribosome footprint density for *ctc* and *yvrE* was much lower compared to their co-transcribed upstream genes. Under σ^B^ induction, additional 5’ ends appeared directly upstream of their coding sequences (Figure 2, red arrows), consistent with the creation of alternative mRNA isoforms from σ^B^-dependent transcription start sites (TSSs, Figure 2 inset). Furthermore, these additional 5’ ends coincide with a sharp increase in ribosome footprint density over the gene bodies.

**Figure 2.**
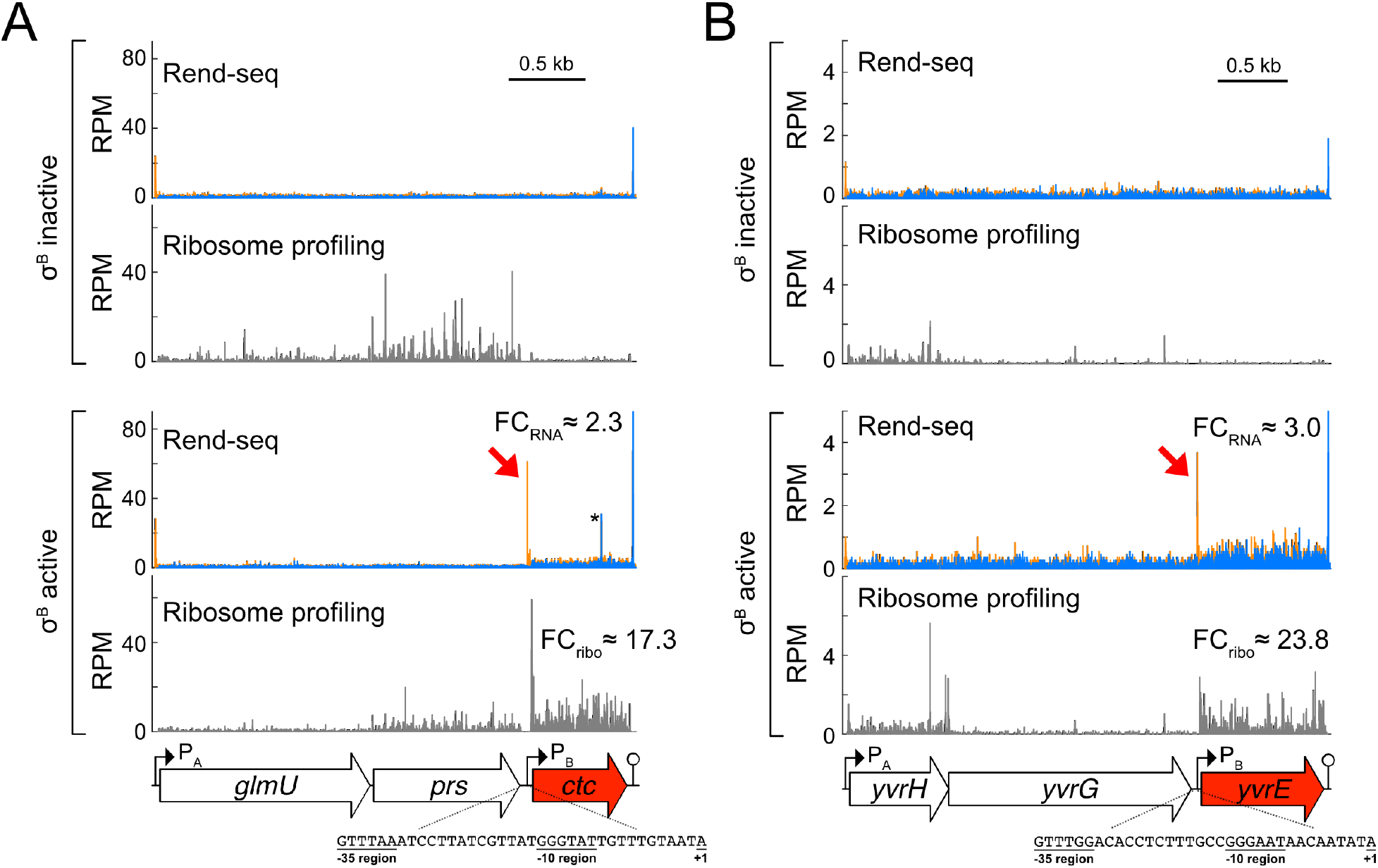
Translationally activated σ^B^ regulon genes display alternative mRNA isoforms. Rend-seq and ribosome profiling data from conditions with inactive/active σ^B^ for the operons containing (**A**) *ctc* and (**B**) yvrE (σ^B^ regulon genes are highlighted in red). Orange and blue bars represent 5’- and 3’-mapped read counts, respectively, and the black scale bars correspond to 0.5 kb. Fold changes (FC) between σ^B^ active and σ^B^ inactive conditions are shown. Rend-seq 5’ ends corresponding to the σ^B^ transcription start sites are marked by red arrows. Putative σ^B^-dependent promoter sequences are listed for each gene (+1 corresponds to the 5’ end of the σ^B^-dependent isoform mapped by Rend-seq). The consensus sequences for the −10 and −35 regions of σ^B^-dependent promoters are GTTTaa and GGG(A/T)A(A/T) (Petersohn et al., 1999). For *ctc* specifically, the additional 5’/3’ peak pair (*) in the σ^B^ active condition corresponds to a spurious RNase A cleavage site that likely occurred post-lysis. See also Figures S1 and S2.

We found that the short, σ^B^-dependent isoforms of the translationally activated genes had significantly elevated translation efficiency compared to the corresponding long, σ^A^-dependent isoforms. By estimating the relative prevalence of short and long isoforms across Rend-seq and ribosome profiling datasets with different levels of σ^B^ induction, we could infer the individual translation efficiency for each isoform (Figure 3A, Figure S3, Methods), hereafter referred to as the isoform-specific translation efficiency. Compared to the σ^A^-dependent isoforms, we found that the TE for the σ^B^ isoform was 3- to 100-fold larger (median = 8.4, Figure 3C). The σ^A^ isoform-specific TEs were all below the median TE across the transcriptome (5/8 in the bottom quartile, Figure 3B), whereas the σ^B^ isoform-specific TEs were all above the median (7/8 in the top quartile). These results indicate that these σ^A^-dependent isoforms are translationally repressed compared to most genes, whereas the σ^B^-dependent isoforms are translationally activated.

**Figure 3.**
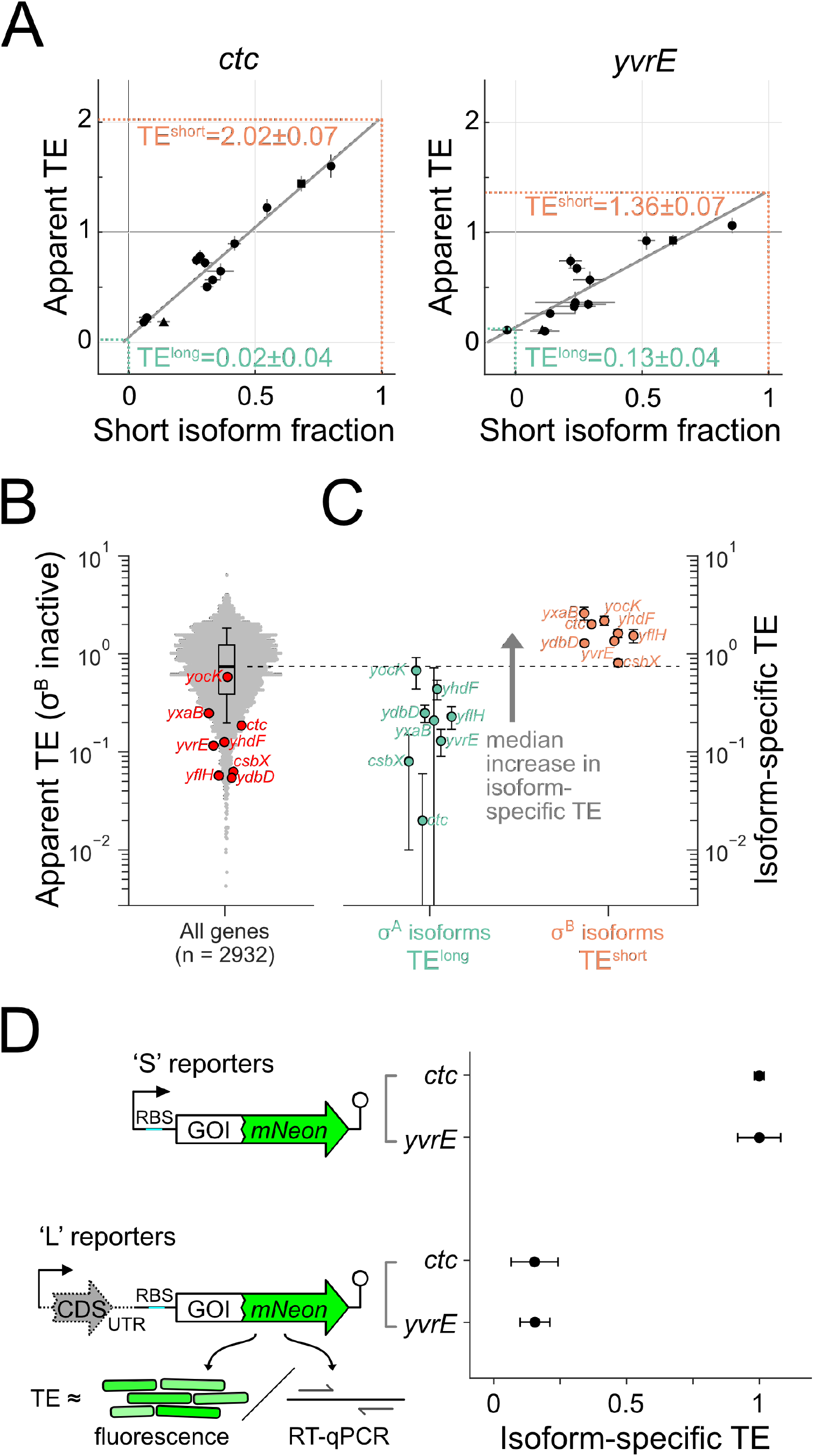
σ^B^-dependent mRNA isoforms have elevated TE. (**A**) Estimation of the isoform-specific TE for the short, σ^B^-dependent and long, σ^A^-dependent isoforms of *ctc* and *yvrE*. Each point is an experimental condition which has a different short isoform fraction and correspondingly different apparent TE (conditions shown in Figure 2 are distinctly marked by a triangle and a square for σ^B^ inactive and active, respectively). Error bars correspond to standard deviations from subsampling bootstraps. The gray lines are linear regressions, whereas the dashed lines indicate estimates of isoform-specific TE calculated from the fits (Methods). Estimated isoform-specific TEs and errors (standard deviations) from a bootstrapped linear fit (Methods) are shown. (**B**) Distribution (beeswarm and boxplot, whiskers corresponding to 10^th^ and 90^th^ percentile) of apparent TE in σ^B^ inactive conditions. Translationally activated σ^B^-regulon genes (subset from Figure 1 for which isoform-specific TE could be estimated, Methods) are marked (red). (**C**) Isoform-specific TE values inferred, with error bars as in (A). (**D**) Fluorescent reporter assay for validating differential TE between isoforms. Protein expression (from fluorescence) and mRNA levels (from reverse-transcription qPCR) were measured for synthetic constructs (left) representing σ^A^-dependent (L) and σB-dependent isoforms (S). Relative isoform-specific TE (right) was calculated by dividing relative protein expression by relative mRNA levels. Errors bars represent the standard deviation for technical replicates (n=3 for fluorescence, n=4 for RT-qPCR). See also Figure S3.

We confirmed that TE was isoform-specific using fluorescent reporter constructs for *ctc* and *yvrE* (Figure 3D). Specifically, we fused the fluorescent protein mNeonGreen to the C-terminal end of each gene. For each fusion protein (*ctc-mNeon, yvrE-mNeon*), two distinct isoform-specific 5’ untranslated region (5’ UTR) variants were placed under the control of an ectopic promoter: 1) a short-isoform variant (S) that included each gene’s native 5’ UTR corresponding to the σ^B^-dependent isoform (as identified by Rend-seq), and 2) a long-isoform variant (L) that included ~100 additional nucleotides in the upstream region, which covers a portion of the coding sequence (CDS) of the upstream gene in the operon. Additionally, a start codon and non-native ribosome binding site (RBS) were inserted directly upstream to enable translation of the truncated upstream CDS in the long-isoform variant. We then quantified the isoform-specific TE for each construct by normalizing relative protein expression (determined from fluorescence, Methods) to relative mRNA levels (from RT-qPCR, Methods). We found that these isoform-specific TEs qualitatively recapitulated our sequencing-based measurements (Figure 3D). Specifically, the isoform-specific TE of the long-isoform constructs was roughly 4- to 6-fold lower than that of the short-isoform constructs, although any further decreases were difficult to quantify due to high background fluorescence. Nevertheless, inclusion of upstream sequence elements was sufficient to produce a large reduction in TE in the absence of the general stress response, which suggests that features in the σ^A^-dependent isoforms can repress translation of the downstream σ^B^ regulon gene. Given the many functions that RNA secondary structure plays in shaping translation in bacteria (Bhattacharyya et al. 2018; Boël et al. 2016; Borujeni et al. 2017; Cambray et al. 2018; Chiaruttini and Guillier 2020; Espah Borujeni and Salis 2016; Goodman et al. 2013; Kudla et al. 2009; Lodish 1968; Li et al. 2014), we aimed to determine if structures in the σ^A^-dependent isoforms could explain the observed impact on translation.

### Extensive secondary structure is associated with translationally repressed, σ^A^-dependent isoforms

To understand the possible role of mRNA secondary structures in setting isoform-specific translation efficiency, we computationally folded for the σ^A^-dependent isoforms of *ctc* and *yvrE*. By mapping the putative Shine-Dalgarno (SD) sequences that recruit ribosome binding (Shine and Dalgarno 1974), onto minimum free energy (MFE) structures (Methods), we found that the majority of bases in the SD sequences were sequestered deep in stable, long-range structures (Figure 4A). Strikingly, in both cases the σ^B^-dependent 5’ ends were located inside the loop of the long RNA stems, such that the short, σ^B^-generated isoforms have their 5’ UTRs entirely liberated from these extended secondary structures. The likelihood of SD sequestration was further supported by calculating the base pairing probability for each position in the SD sequences, which revealed that the majority of positions were predicted to be paired across the full thermodynamic ensemble (base-pairing probability ≈1). Given that SD sequences facilitate ribosome recruitment to mRNA to initiate translation, we expected that the presence of extensive secondary structure at and around these elements in the σ^A^-dependent isoforms could plausibly repress translation of the downstream σ^B^ regulon gene. However, numerous factors in the cellular microenvironment affect the folding dynamics of RNAs, yielding *in vivo* structures that can differ substantially from their *in silico* counterparts (Mustoe et al. 2018; Rouskin et al. 2014; Spitale et al. 2015; Burkhardt et al. 2017). Accordingly, we decided to experimentally validate these computationally predicted structures for the σ^A^-dependent isoforms of *ctc* and *yvrE*.

**Figure 4.**
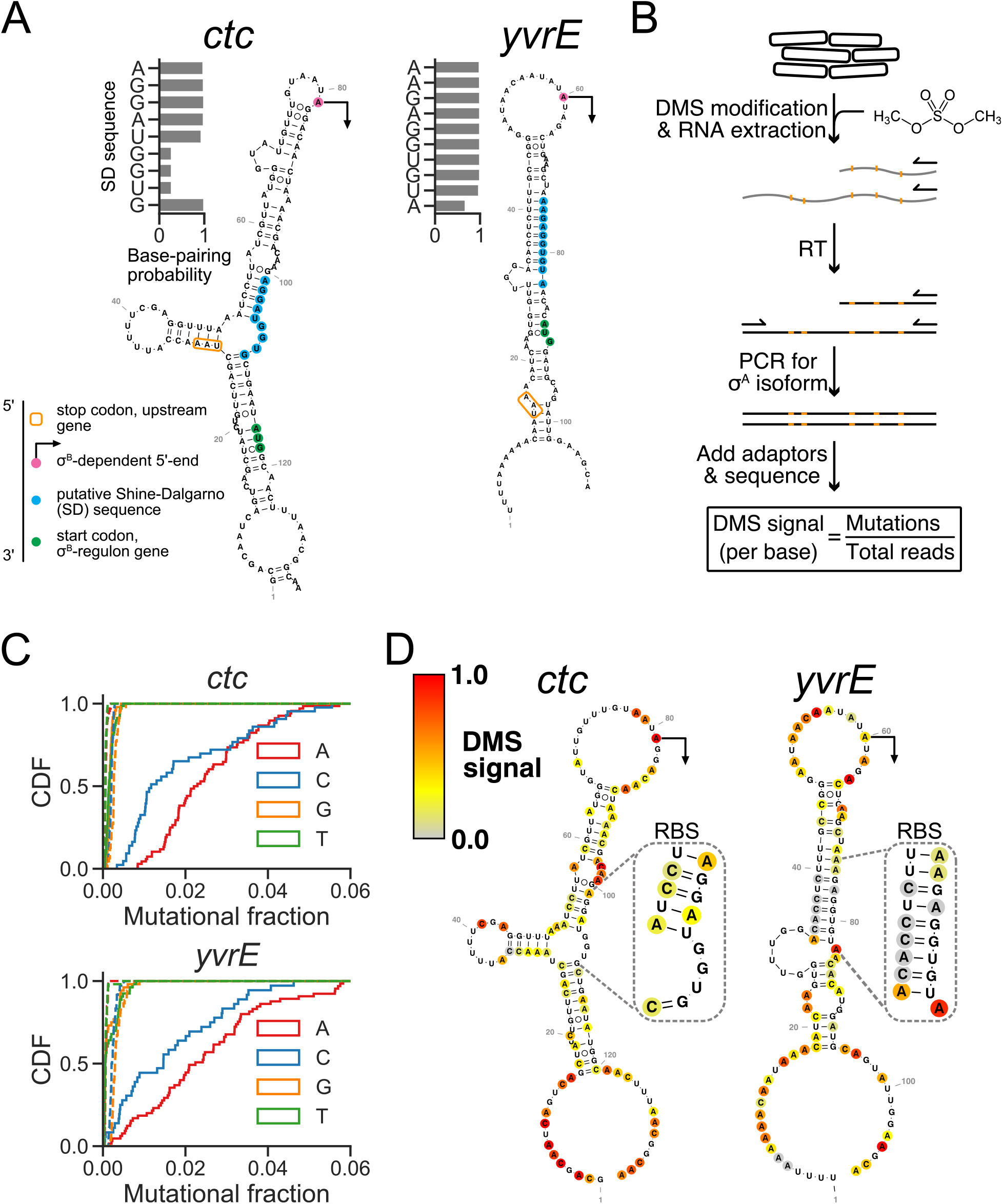
σ^A^-dependent mRNA isoforms have extended secondary structures *in vivo*. (**A**) Minimum free energy (MFE) structures of the σ^A^-dependent isoforms of *ctc* and *yvrE* near the ribosome binding site. The transcription start sites of σ^B^-dependent isoforms (indicated with arrows), putative Shine-Dalgarno (SD) sequences, and start codons are highlighted in magenta, blue, and green, respectively. The stop codon of the upstream gene in the operon is indicated with an orange box. Computationally-determined base-pairing probabilities for individual bases in the SD sequences are shown beside each structure. (**B**) DMS-MaPseq workflow for *in vivo* RNA structure determination of σ^A^-dependent isoforms. (**C**) Cumulative distributions of the per-base mutational fractions for the σ^A^-dependent isoforms of *ctc* and *yvrE*. Solid and dashed lines indicate conditions with and without DMS treatment. (**D**) DMS-constrained MFE structures of representative transcripts for σ^A^-dependent isoforms of *ctc* and *yvrE* colored by normalized DMS-MaPseq mutation rate (DMS signal), where values correspond to increased base accessibility. Structured regions containing putative SD sequences are magnified.

We employed the RNA structure probing method DMS-MaPseq to quantify mRNA structures *in vivo*. This technique involves treating RNA with the methylating agent dimethyl sulfate (DMS) to modify the base-pairing faces of accessible adenine and cytosine nucleobases. These modifications are subsequently encoded as mutations during reverse transcription using a specialized thermostable group II intron reverse transcriptase, generating a mutational signal that is detectable using high-throughput sequencing and has been shown to correlate with base accessibility (Tomezsko et al. 2020; Zubradt et al. 2016). We used a targeted version of DMS-MaPseq to specifically reverse transcribe and amplify the predicted structural region in the σ^A^-dependent isoforms of *ctc* and *yvrE* following DMS treatment *in vivo* (Figure 4B). After sequencing these amplicons, we examined the per-base mutational fractions against a control without DMS treatment and confirmed that DMS induced a characteristic signal at amino bases (Figure 4C).

We refolded the σ^A^-dependent isoforms of *ctc* and *yvrE* using DMS signal as a constraint (Methods) and found strong agreement with the earlier MFE structures (Figure 4D). In particular, the regions containing the SD sequences were indeed highly structured *in vivo* and thus less accessible to the translation machinery. Additionally, these structured regions were robust to the folding window size (Methods). These extended structures that occlude the ribosome binding sites are consistent with the repressed translation of the long, σ^A^-dependent isoforms.

After validating the computationally predicted secondary structures by DMS-MaPseq, we extended our computational analysis to additional translationally activated σ^B^ regulon genes and found a consistent pattern of characteristic structures in the σ^A^-dependent isoforms that sequester the sequence elements required for translation initiation (Figure 5). Similar to *ctc* and *yvrE*, the remaining 6 genes for which we estimated isoform-specific TE all displayed MFE structures with the SD sequences located in extended stem-loops, and base pairing probabilities indicated that the SD sequences were predominantly paired. These results suggest that these other σ^A^-dependent long isoforms are also translationally repressed by extensive secondary structures, like the orthogonally validated instances of *ctc* and *yvrE*.

**Figure 5.**
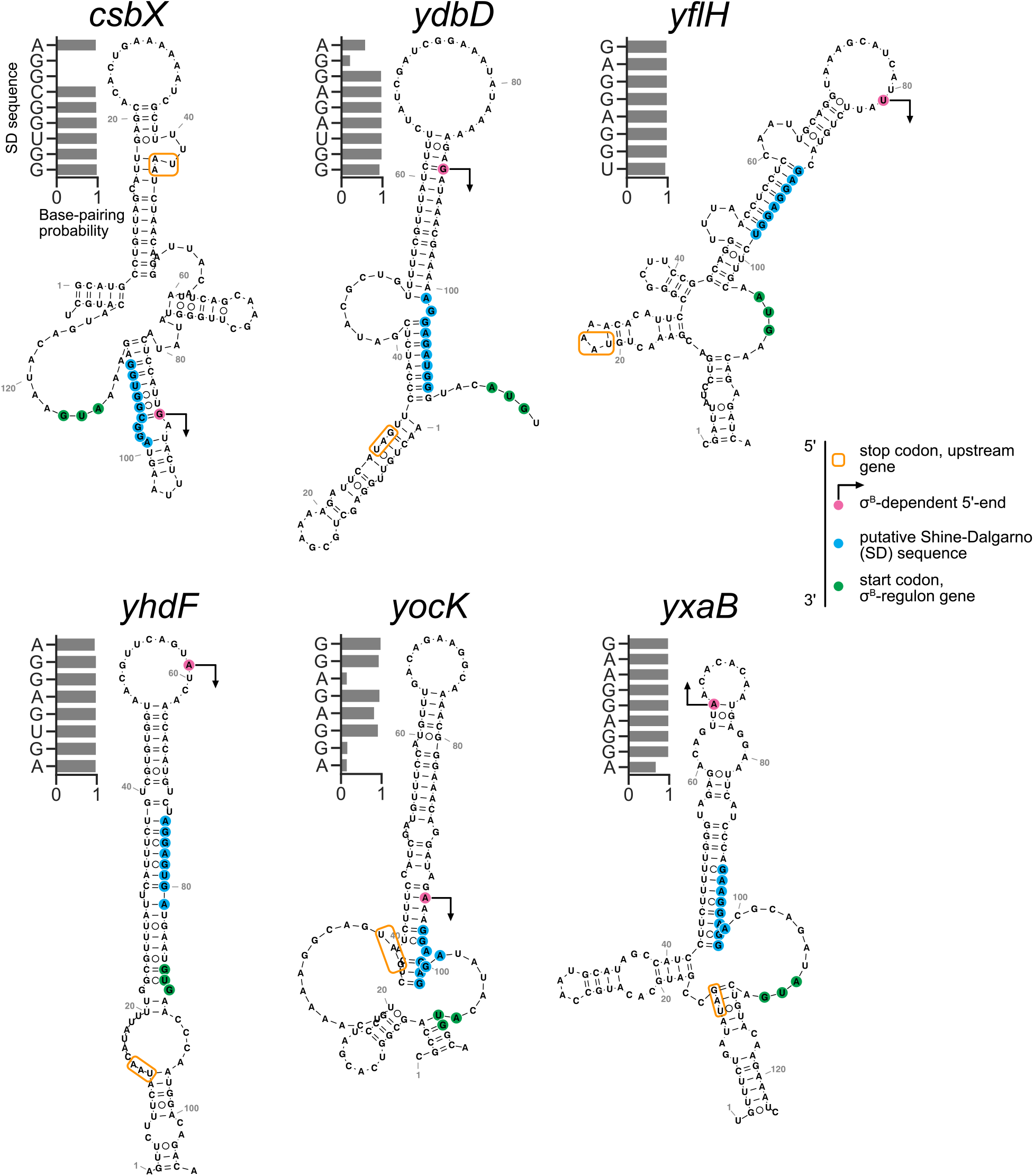
Long-range mRNA secondary structures in σ^A^-dependent isoforms sequester sequence elements necessary for translation. MFE structures of transcripts for σ^A^-dependent isoforms of other translationally activated σ^B^ regulon genes. The transcription start sites of σ^B^-dependent isoforms (indicated with arrows), putative Shine-Dalgarno (SD) sequences, and start codons are highlighted in magenta, blue, and green, respectively. The stop codon of the upstream gene in the operon is indicated with an orange box. Computationally-determined base-pairing probabilities for individual bases in the SD sequences are shown beside each structure.

### Internal σ^B^ promoters liberate mRNA secondary structure and activate translation

In contrast to being repressed in the σ^A^-dependent isoforms, genes in the short, σ^B^-dependent isoforms had above-normal levels of translation (Figure 3C). The single-nucleotide resolution afforded by Rend-seq data revealed a common feature among this group of genes: the TSSs of the σ^B^-dependent isoforms were located within the extended secondary structure, often inside the loop region or in the downstream stem (Figure 4A, Figure 5, magenta and arrow). Therefore, σ^B^-driven transcription generates isoforms with 5’ UTRs that lack the upstream portion of the stem sequestering the SD sequence in the long, σ^A^-dependent isoforms, thereby freeing up the ribosome binding site for efficient translation initiation.

The prevalence of this regulation suggests an alternative configuration for σ^B^-dependent gene expression that does not entirely rely on its canonical role as acting at the transcriptional level. In this operonic architecture, σ^A^-driven promoters produce long, polycistronic mRNAs containing stable structures that impede translation initiation for σ^B^ regulon genes located at the ends of these transcripts (Figure 6). When activated by stress, however, σ^B^ initiates transcription from alternative promoters directly upstream of its regulon genes, bypassing the inhibitory secondary structures and thereby promoting ribosome binding on these shorter mRNAs. The resulting increase in protein expression predominantly arises from a greater ribosome flux on these transcripts, demonstrating a novel function for σ^B^ in regulating gene expression in a simultaneous transcriptional-translational induction.

**Figure 6.**
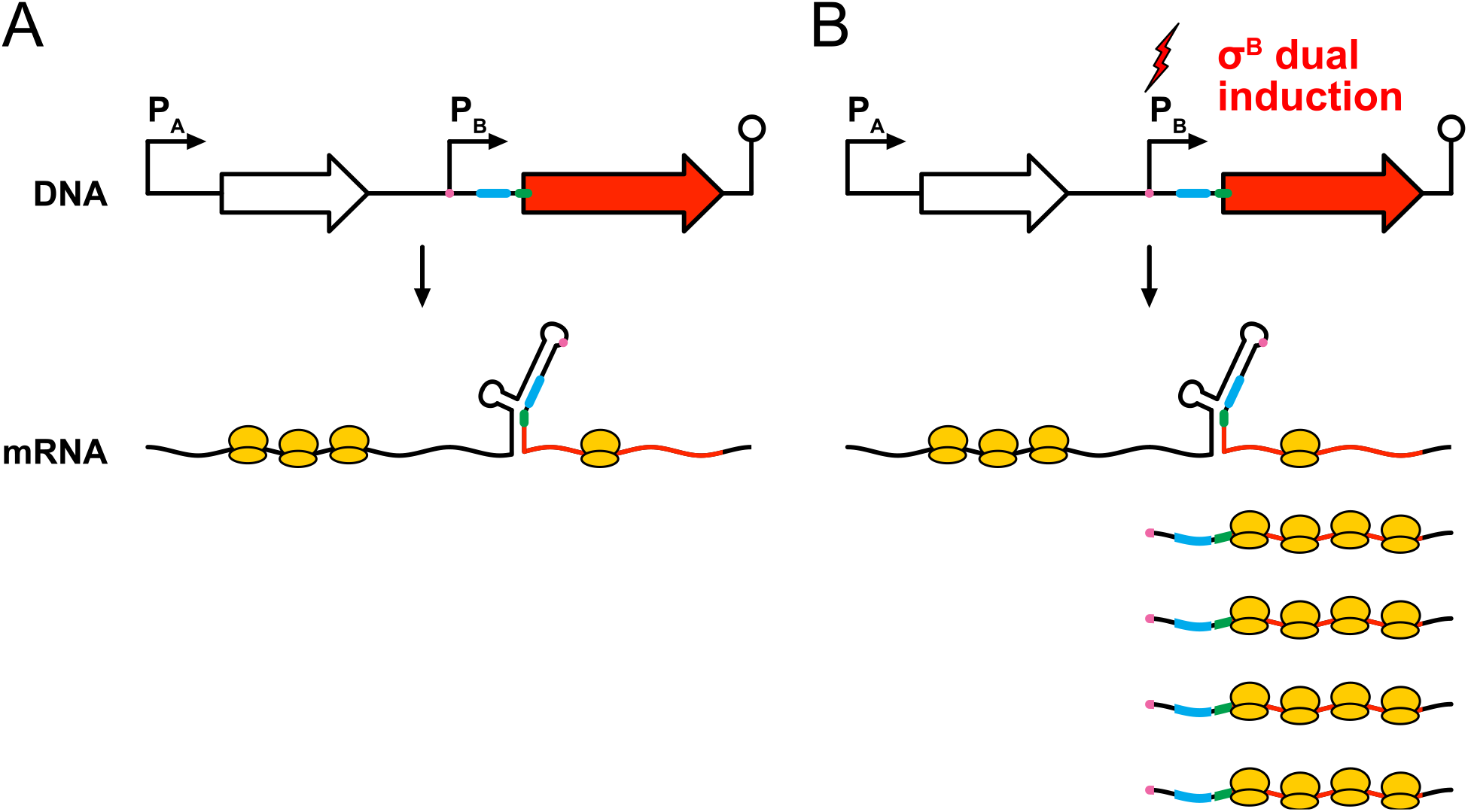
Model for σ^B^-dependent translational activation. Schematic of a polycistronic operon containing a σ^A^-dependent promoter (P_A_), σ^B^-dependent promoter (P_B_), coding sequences, and a terminator. (**A**) In the absence of σ^B^, transcription from PA produces a polycistronic mRNA molecule containing secondary structures that translationally repress the σ^B^-dependent open reading frame (red) by sequestering its Shine-Dalgarno sequence (blue) and start codon (green). (**B**) P_B_ becomes transcriptionally active upon σ^B^ induction, generating an mRNA isoform with an alternative transcription start site (magenta). Without the sequences necessary to form stable secondary structures, these transcripts can recruit ribosomes more efficiently to facilitate greater protein expression.

## DISCUSSION

Bacterial sigma factors have long been studied as quintessential examples of gene regulation. Mechanistically, their direct effects on transcription initiation are well-understood (Paget 2015). We expand this view by demonstrating that the alternative sigma factor σ^B^ in *B. subtilis* can also influence translation initiation for several of its regulon genes. Translation activation is accomplished by modulating isoform-specific RNA secondary structures that normally impede translation initiation. This multifunctional control of transcription and translation by a single trans-acting factor serves as a strategy to enable massive upregulation of gene expression under specific cellular conditions.

The RNA secondary structures that impede translation in the long, σ^A^-dependent isoforms often include regions of the upstream open reading frames (ORFs), raising questions about whether ribosomes translating the upstream ORFs may perturb the formation of the inhibitory secondary structures. Ribosomes are known to unwind structured regions of RNA as they elongate over coding sequences (Takyar et al. 2005; Wen et al. 2008). We observed that the stop codon of the upstream gene in the operon was typically located within the large stem-loop (Figure 4, Figure 5). This places ribosomes in proximity to the critical structural elements if the upstream message is actively translated. However, the results from our fluorescent reporter assay show that this configuration is not capable of fully restoring translation for either *ctc* or*yvrE*, despite the upstream gene being driven by an exogenous ribosome binding site with the consensus SD sequence. These data suggest that translation of the upstream gene is insufficient to fully derepress downstream genes, presumably because the ribosome footprint does not extend sufficiently downstream to disrupt RNA structure, or possibly due to rapid refolding of secondary structures after ribosomes pass through.

What is the utility of this regulatory strategy? From an evolutionary perspective, it seems counterintuitive for these genes to be found within larger operons despite being lowly translated. We could instead imagine a transcription terminator evolving in the region between the upstream genes in the operon and the σ^B^-dependent TSS, which would ensure that the σ^B^ regulon gene is only induced upon activation of the general stress response. One potential explanation for multifunctional regulation is to allow fine-tuned expression of some σ^B^ regulon genes during non-stress conditions. On the one hand, this transcript architecture enables these genes to be transcribed during exponential growth. On the other hand, translation may have been selected against in the same condition to avoid fitness defects from overexpression. In this case, the observed basal expression from the σ^A^-driven isoforms would be sufficient for their functions during non-stress conditions.

Another possible explanation for this regulatory strategy could be that small amounts of these proteins are necessary for coping with general stress during transitional periods where σ^B^ has already been activated but synthesis of general stress proteins is still ongoing. A fitness benefit would be challenging to identify except in specific conditions where the cell relies on one of these particular σ^B^ regulon genes for survival. Indeed, extensive phenotyping of σ^B^-regulon member deletions under varied stresses has demonstrated the limited impact of individual proteins on cell fitness (Höper et al. 2005). Identifying the exact stress conditions in which this regulatory strategy confers a fitness advantage constitutes an interesting future direction.

Regardless of the function of σ^B^-dependent translational activation, our discovery and characterization of sigma factor-mediated dual induction (Figure 6) expands our view of the regulatory roles of sigma factors and reveals an intriguing principle of bacterial genome organization that could be further investigated in similar organisms.

## MATERIALS AND METHODS

### Strains and strain construction

Strains used to generate new data in this study are listed in Table 1. Strains pertaining to matched Rend-seq and ribosome profiling datasets (from GEO accession GSE162169) are listed in Table S1.

**TABLE 1.**
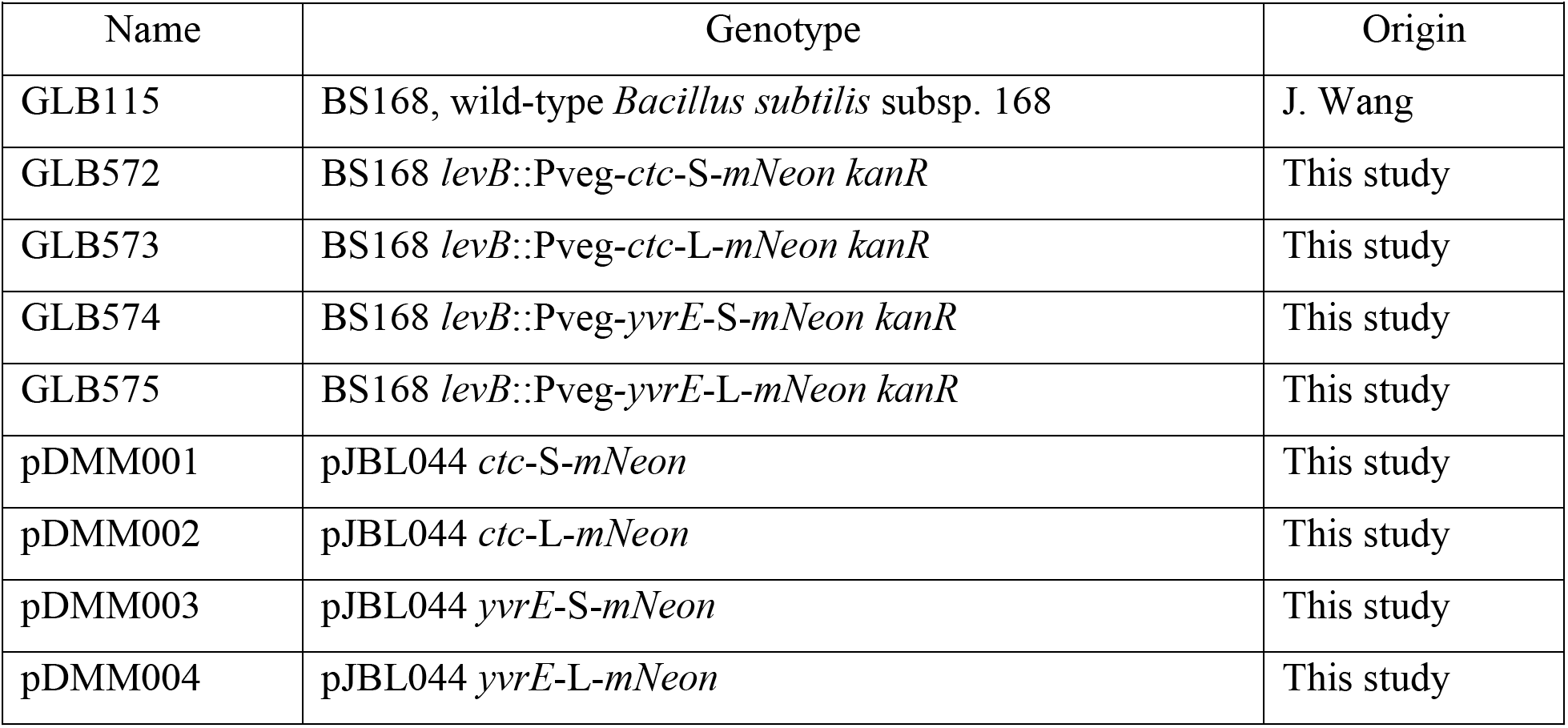
Strains and plasmids used in this study.

To construct the strains for the fluorescent reporter assay, the genes *ctc* and *yvrE* (with variable upstream regions) were fused to the fluorescent protein mNeonGreen with a C-terminal linker and cloned into pJBL044 under the constitutive promoter Pveg using Gibson assembly (New England Biolabs). The original pJBL044 plasmid was constructed using isothermal assembly from a fragment of pDR160 (Bose and Grossman 2011), a *kanR* cassette (Guérout-Fleury et al. 1995), *levB* homology regions, the Pveg promoter, and the strong *efp* terminator. The assembled plasmids were transformed into *Mix and Go! E. coli* DH5 Alpha Competent Cells (Zymo Research) per the manufacturer’s instructions and isolated using a QIAprep Spin Miniprep Kit (QIAGEN). The fusion constructs were then integrated into BS168 at the *levB* locus using standard cloning techniques (Harwood, C R and Cutting 1990), and successful recombinants were verified by colony PCR. All plasmids and recombinants (see Table 1) were further validated by Sanger sequencing (Quintara Biosciences).

### Growth conditions

Unless indicated otherwise, all strains were grown at 37°C with shaking (250 rpm) in LB supplemented with carbenicillin (100 μg/mL for *E. coli*) and/or kanamycin (50 μg/mL for *E. coli*, 5 μg/mL for *B. subtilis*) when appropriate. For overnight cultures, LB liquid media was inoculated with single colonies from LB agar plates.

For matched Rend-seq/ribosome profiling datasets, strains were grown in LB or conditioned MCC medium (Parker et al. 2020; Lalanne et al. 2020) with various inducer (xylose, IPTG) concentrations (see Table S1). For these datasets, cells were grown in exponential phase for at least 10 doublings before harvesting at OD_600_≈0.3.

### Existing Rend-seq and ribosome profiling datasets

Matched Rend-seq and ribosome profiling datasets used to identify genes with increased TE (Figure 1) and to estimate the short isoform fraction and corresponding apparent TE (Figure 3, Figure S3) were obtained from GEO accession GSE162169 (Lalanne et al. 2020). These datasets display a range of σ^B^ activation due to a diverse set of genetic modifications and growth media. In particular, we previously identified that tuning the expression of translation termination factors RF2 and PrmC activate σ^B^ to varying degrees (Lalanne et al. 2020). For example, the σ^B^ active data presented in Figures 1 and 2 correspond to a CRISPRi knockdown of RF2, while σ^B^ inactive corresponds to wild-type. Importantly, although it is possible that different RF2 levels could affect translation initiation (and therefore TE) of genes (Lalanne et al. 2020), none of the genes that show a substantial increase in TE (Figure 1) have a UGA stop codon or are co-transcribed with a gene ending with UGA stop (UGA being the stop codon cognate to RF2). Hence, the molecular causes of σ^B^ activation are distinct and independent from the mechanisms leading to translational activation characterized here.

### Quantification of mRNA level, ribosome footprint density, and translation efficiency

From pile-up files (.wig format), the mRNA level corresponding to a gene was quantified as the 1% winsorized average read density for 3’-end mapped Rend-seq reads across the body of the gene, excluding a 40 nt region the start and end of the gene (start+40 nt to end-40 nt for averaging). Ribosome footprint read density was similarly calculated (1% winsorized density from start+40 nt to end-40 nt). Read densities were then normalized to rpkm (reads per kilobase per million reads mapped) using the total number of reads mapping to non-rRNA or tRNAs. For all genes, bootstrap (randomly sampling from the distribution of read counts per position across the body of the gene and calculating the corresponding resampled density and downstream quantities) was used as a measure of technical and read count variability. Error bars in Figures 3A and S3 correspond to the standard deviation across bootstrap subsamplings. Large error bars correspond to large counting noise (regions with few reads mapped). The translation efficiency of each gene was calculated as the ribosome profiling rpkm divided by the Rend-seq rpkm. Only genes with >50 reads mapped were considered to identify candidates with substantially elevated TE (Figure 1).

### Determination of isoform-specific TE

To estimate the isoform-specific TE for particular genes, we assume that each individual mRNA isoform has a distinct TE, and that the total ribosome footprint density for a gene with multiple mRNA isoforms is equal to the sum of the isoform-specific TEs weighted by the mRNA abundance of each isoform.

Specifically, consider a two-gene operon with a long isoform that includes both gene 1 and gene 2 as well as a short isoform that contains gene 2 exclusively (schematically illustrated in Figure S3A). Denote overall mRNA level for genes 1 and 2 by *m*_1_ and *m*_2_, and overall ribosome footprint density *r*_1_ and *r*_2_ for the two genes respectively. Further, let *m_short_, m_long_* be the level of the short and long isoform respectively, and *TE*^2,*short*^, *TE*^2,*long*^ the corresponding isoform-specific TE. Note that the overall mRNA level for genes 1 and 2 are related to isoform mRNA levels by: *m*_1_ = *m_long_* and *m*_2_ = *m_short_* + *m_lon_*. Hence, from the total mRNA level for both genes, we can infer the isoform mRNA levels: *m_long_* = *m*_1_, and *m_short_* = *m*_2_ – *m*_1_.

By assumption, for the ribosome density on gene 2: *r*_2_ = *m_short_TE*^2,*short*^ + *m_long_TE*^2,*long*^. For the apparent TE of gene 2, we thus have: 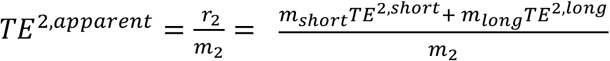. Reorganizing the equation leads to: *TE*^2,*apparent*^ = *f_short_TE*^2, *short*^ + (1 – *f_short_*)*TE*^2, *long*^, were we have defined the short isoform mRNA fraction for gene 2 as 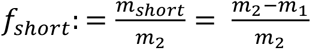. We note that for genes in conditions with little to no short isoform expression, the estimated short isoform fraction may be negative as a result of the technical variability in coverage.

Using RNA-seq data, *f_short_* can be estimated from the mRNA levels on both genes as shown above as 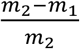. Using ribosome profiling data from a matched sample, the apparent TE on gene 2, *TE*^2, *apparent*^, can be estimated as 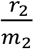. If our assumption of isoform-specific TE linearly contributing to overall ribosome density on gene 2 is valid, then a plot of *TE*^2,*apparent*^ vs. *f_short_* across samples with variable induction of the short isoform should display a linear relationship, with a y-intercept at *f_short_* = 0 of *TE*^2,*long*^ and a y-intercept at *f_short_* = 1 of *TE*^2,*short*^ as seen in Figures 3A and S3B.

To increase the precision of the determination of the short and long isoform mRNA levels, genomic regions used to quantify mRNA levels were extended beyond gene bodies using manually curated transcript boundaries determined by Rend-seq. mRNA levels and ribosome footprint densities were calculated as the average read densities across these regions in Rend-seq and ribosome profiling data, respectively.

To determine the uncertainty on estimated isoform-specific TEs, linear regressions were performed on bootstrap resampling estimates for the short isoform fractions and apparent TEs. Each bootstrap regression provided an estimated TE^long^ and TE^short^. The error bars for these quantities (Figure 3A, Figure 3C, Figure S3B) were taken as the standard deviations of these bootstrap estimates.

For the genes that do not belong to the group with the characteristic long, σ^A^-dependent isoforms and short, σ^B^-dependent isoforms (Figure S2), their alternative promoters are too close to allow proper quantification of isoform-specific abundances. These were thus excluded from the above analyses.

### Fluorescent reporter assay

For the fluorescence reporter assay, the strains GLB115, GLB572, GLB573, GLB574, and GLB575 were grown to OD_600_≈1-2 and then back-diluted 200-fold into fresh media. Three technical replicates per culture were grown at 37°C for 12 hr in a BioTek Synergy H1 microplate reader, and absorbance (600 nm) and fluorescence intensity (EX 485/20 nm, EM 520/20 nm) were measured every 5 min. Fluorescence was normalized by absorbance at each time point, and any background signal from cellular/media autofluorescence was removed by subtracting the mean normalized fluorescence values of the wild-type BS168 replicates. These quantities were then converted to relative values by normalizing proportionally to the signal for the S variants.

For RT-qPCR, overnight cultures of the same strains were back-diluted to OD_600_≈2×10^-4^ and regrown for roughly 10 generations. At OD_600_≈0.3, 5 mL of cells were harvested and mixed with 5 mL of chilled methanol, spun down at 4°C for 10 min, and frozen at −80°C after removing the supernatant. Thawed cell pellets were treated with 100 μL of 10 mg/mL lysozyme in TE, and total RNA was extracted using a RNeasy Mini Kit (QIAGEN). DNA was removed using TURBO DNase (Thermo Fisher Scientific), and RNA was purified using isopropanol precipitation. Reverse transcription was performed using Random Hexamer Primer (Thermo Fisher Scientific) and M-MuLV Reverse Transcriptase (New England Biolabs) per the manufacturer’s instructions. RNA levels were measured on a Roche LightCycler 480 Real-Time PCR system using two primer sets for *mNeon* and one primer set each for the loading controls *gyrA* and *sigA (mNeon* F1, *mNeon* R1, *mNeon* F2, *mNeon* R2, *gyrA* F, *gyrA* R, *sigA* F, *sigA* R, see Table 2). The fold change in *mNeon* RNA levels relative to the S strains was calculated by taking the average of three technical replicates across each combination of primer sets (*mNeon1/gyrA, mNeon1/sigA, mNeon2/gyrA, mNeon2/sigA*).

**TABLE 2.**
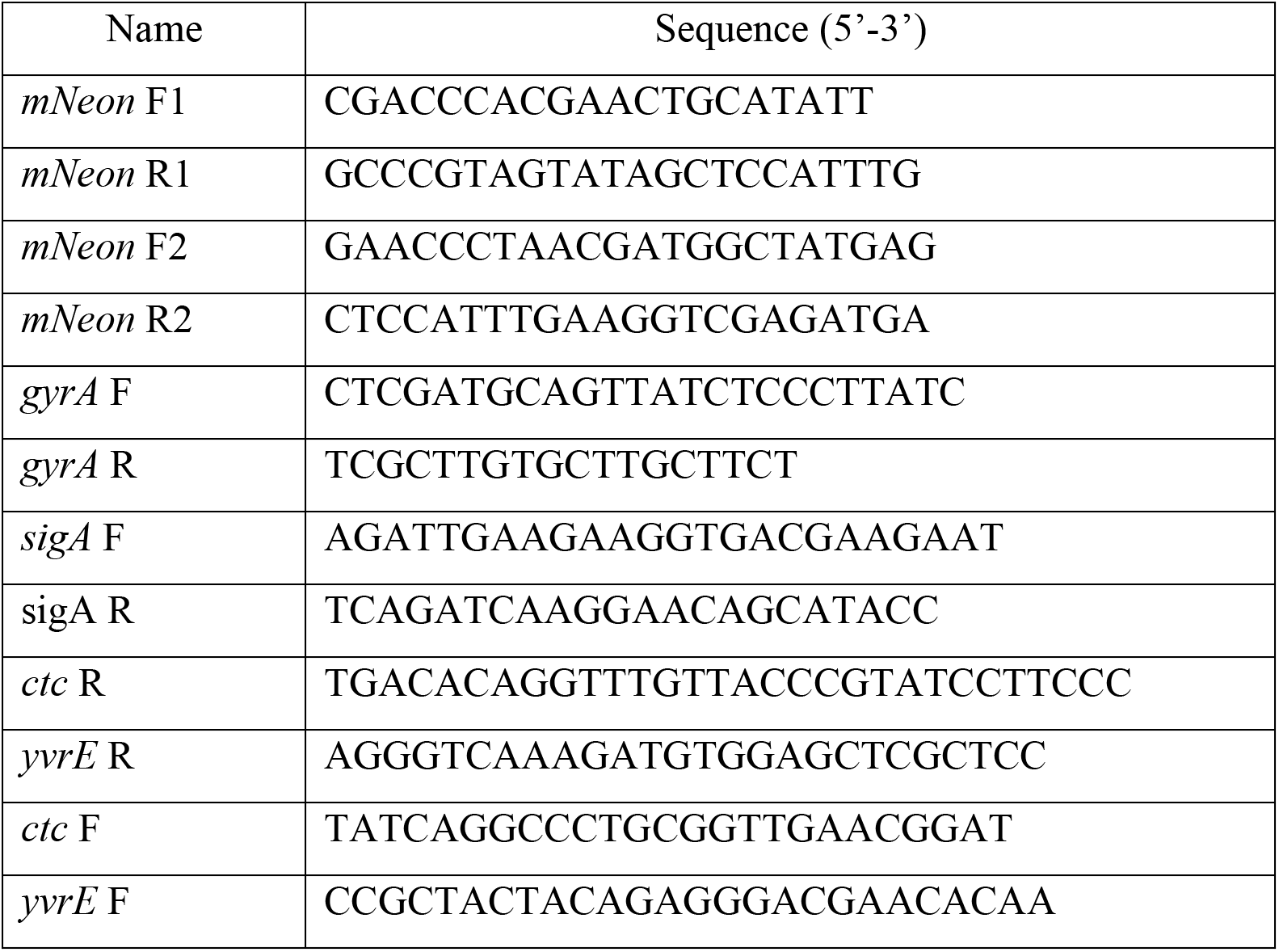
Oligos used in this study.

Isoform-specific TE was subsequently calculated by normalizing mean relative fluorescence by mean fold change in *mNeon* RNA levels, and the standard deviation was propagated from each measurement type.

### RNA secondary structure prediction

Minimum free energy (MFE) structures were predicted using the RNAfold program of the ViennaRNA Package (Lorenz et al. 2011) with default parameters. Base-pairing probabilities were determined by constraining each position in a sequence individually as unpaired and then calculating the partition function from the ensemble free energy computed by RNAfold. The probability of each position being unpaired was calculated by dividing the partition function for the constrained sequence by the partition function for an unconstrained sequence, and the base pairing probabilities were simply the probabilities of the complements. Putative Shine-Dalgarno (SD) sequences were identified as the region upstream of the start codon that forms the strongest duplex with the anti-Shine-Dalgarno (aSD, 5’-TCACCTCCT-3’) sequence in the 16S ribosomal RNA. RNA secondary structures determined using RNAfold were visualized using VARNA v3.93 (Visualization Applet for RNA) (Darty et al. 2009). The structures sequestering the ribosome binding sites shown in Figures 4 and 5 were confirmed to be robust to the specific regions computationally folded, both at the level of secondary structure and base-pairing probabilities of the SD sequences.

### DMS-MaPseq

*In vivo* DMS treatment was performed as previously described (Burkhardt et al. 2017; Zubradt et al. 2016). Specifically, an overnight culture of BS168 was split two ways and back-diluted to OD_600_≈2×10^-4^. Following regrowth to OD_600_≈0.2, 15 mL of each culture was incubated at 37°C for 2 min with shaking (1000 rpm) after treating one with 750 μL of dimethyl sulfate (DMS, ~5% final concentration). The reaction was stopped by adding 30 mL of chilled stop solution (30% β-mercaptoethanol, 25% isoamyl alcohol) to each sample, after which they were immediately transferred to ice and spun down at 4°C for 8 min. The cell pellets were washed with 8 mL of chilled wash solution (30% β-mercaptoethanol), resuspended in residual wash solution, and frozen at −80°C. Thawed cell pellets were treated with 100 μL of 10 mg/mL lysozyme in TE, and total RNA lysis buffer (10 mM EDTA, 50 mM sodium acetate) was added to 650 μL. Total RNA was extracted using hot acid-phenol:chloroform and isopropanol precipitation.

For library preparation, established protocols (Tomezsko et al. 2020; Zubradt et al. 2016) were again followed. DNA was removed using TURBO DNase, and RNA >200 nt was purified using an RNA Clean & Concentrator-5 Kit per the manufacturer’s instructions (Zymo Research). Ribosomal RNA was depleted using a MICROBExpress Bacterial mRNA Enrichment Kit (Thermo Fisher Scientific), and RNA >200 nt was again purified using an RNA Clean & Concentrator-5 Kit. Reverse transcription was performed at 64°C for 90 min using 70 ng of RNA from each sample and TGIRT-III (Ingex). The RT primers were specific to each gene (*ctc* R, *yvrE* R, see Table 2). The RT reaction was treated with 1 μl RNase H (New England Biolabs) and incubated at 37°C for 20 min to remove RNA. Roughly 1/10 of the resulting volume was used as template for a two-step PCR amplification with Phusion High-Fidelity DNA Polymerase (New England Biolabs) per the manufacturer’s specifications, which was run for 15-25 cycles with the RT primer serving as the reverse primer (*ctc* F, *yvrE* F, see Table 2). PCR products (~240-290 bp) were purified by gel extraction on an 8% TBE polyacrylamide gel (Thermo Fisher Scientific) and isopropanol precipitation. Samples with particularly low dsDNA concentrations (as measured on an Invitrogen Qubit 4 Fluorometer) were reamplified for 7-20 additional cycles and purified in the same manner. After adding adapters via PCR, the libraries were sequenced on an Illumina MiSeq (2 × 250 nt reads).

To determine the DMS signal, FASTQ files were processed and analyzed using the DREEM (Detection of RNA folding Ensembles using Expectation-Maximization clustering) pipeline with the ‘--fastq’ and ‘--struct’ options (Tomezsko et al. 2020). In brief, paired-end reads were filtered for quality and trimmed using FASTQC v.0.11.8 and TrimGalore 0.4.1, respectively. Reads were aligned to target sequences in the reference genome NC_000964.3 from the NCBI RefSeq database using Bowtie2 2.3.4.1 with the options ‘--local --no-unal --no-discordant --no-mixed −X 1000 −L 12’. Mapped reads were represented as bit vectors and clustered by their mutational signatures using the DREEM algorithm with standard parameters (Tomezsko et al. 2020). Per-base mutational fractions were initially quantified using the population-average fraction of mismatches and deletions. Following expectation-maximization (EM) clustering, the DMS reactivity was taken as the mutation rates of the bases in the cluster K=1. After normalizing to the median of the top 5% of positions (with the upper limit set to 1.0), the DMS signal was used as a folding constraint for predicting RNA secondary structures with the program RNAstructure v.6.0.1 (Reuter and Mathews 2010) Additionally, the folding windows were expanded symmetrically by 50, 100, 150, and 200-nt in either direction to assess the robustness of the predicted folds. RNA secondary structures were again visualized using VARNA v3.93 (Darty et al. 2009). The sequencing datasets for DMS-MaPseq are available online using the GEO accession GSE168393.

## SUPPLEMENTAL MATERIAL

Supplemental material (Figures S1-S3, Table S1) is available for this article.

## ACKNOWLEDGEMENTS

We thank members of the Li lab and the Rouskin lab for critical discussions. We thank S. McKeithen-Mead for providing the *mNeon* template. This research was supported by NIH grant R35GM124732, the NSF CAREER Award, the Smith Odyssey Award, the Pew Biomedical Scholars Program, a Sloan Research Fellowship, the Searle Scholars Program, the Smith Family Award for Excellence in Biomedical Research; NSERC doctoral Fellowship and HHMI International Student Research Fellowship (to J.-B.L.).

## REFERENCES

Bastet L, Turcotte P, Wade JT, Lafontaine DA. 2018. Maestro of regulation: Riboswitches orchestrate gene expression at the levels of translation, transcription and mRNA decay. RNA Biol.

Bhattacharyya S, Jacobs WM, Adkar B V., Yan J, Zhang W, Shakhnovich EI. 2018. Accessibility of the Shine-Dalgarno Sequence Dictates N-Terminal Codon Bias in E. coli. Mol Cell 70.

Boël G, Letso R, Neely H, Price N, Wong K, Su M, Luff JD, Valecha M, Hunt JF, Everett JK, et al. 2016. Codon influence on protein expression in E. coli correlates with mRNA levels. Nature. http://dx.doi.org/10.1038/nature16509.

Borujeni AE, Cetnar D, Farasat I, Smith A, Lundgren N, Salis HM. 2017. Precise quantification of translation inhibition by mRNA structures that overlap with the ribosomal footprint in N-terminal coding sequences. Nucleic Acids Res 45.

Bose B, Grossman AD. 2011. Regulation of horizontal gene transfer in Bacillus subtilis by activation of a conserved site-specific protease. J Bacteriol 193: 22–29.

Breaker RR. 2018. Riboswitches and translation control. Cold Spring Harb Perspect Biol.

Burkhardt DH, Rouskin S, Zhang Y, Li GW, Weissman JS, Gross CA. 2017. Operon mRNAs are organized into ORF-centric structures that predict translation efficiency. Elife.

Cambray G, Guimaraes JC, Arkin AP. 2018. Evaluation of 244,000 synthetic sequences reveals design principles to optimize translation in escherichia coli. Nat Biotechnol 36: 1005.

Chauvier A, Picard-Jean F, Berger-Dancause JC, Bastet L, Naghdi MR, Dubé A, Turcotte P, Perreault J, Lafontaine DA. 2017. Transcriptional pausing at the translation start site operates as a critical checkpoint for riboswitch regulation. Nat Commun.

Chiaruttini C, Guillier M. 2020. On the role of mRNA secondary structure in bacterial translation. Wiley Interdiscip Rev RNA 11.

Darty K, Denise A, Ponty Y. 2009. VARNA: Interactive drawing and editing of the RNA secondary structure. Bioinformatics.

Espah Borujeni A, Salis HM. 2016. Translation Initiation is Controlled by RNA Folding Kinetics via a Ribosome Drafting Mechanism. J Am Chem Soc jacs.6b01453. http://pubs.acs.org/doi/abs/10.1021/jacs.6b01453.

Goodman DB, Church GM, Kosuri S. 2013. Causes and effects of N-terminal codon bias in bacterial genes. Science (80-) 342.

Guérout-Fleury AM, Shazand K, Frandsen N, Stragier P. 1995. Antibiotic-resistance cassettes for Bacillus subtilis. Gene 167: 335–336.

Haldenwang WG. 1995. The sigma factors of Bacillus subtilis. Microbiol Rev.

Haldenwang WG, Losick R. 1979. A modified RNA polymerase transcribes a cloned gene under sporulation control in Bacillus subtilis. Nature 282.

Harwood, C R and Cutting SM. 1990. Molecular Biological methods for Bacillus. John Wiley.

Hecker M, Pané-Farré J, Uwe V. 2007. SigB-Dependent General Stress Response in *Bacillus subtilis* and Related Gram-Positive Bacteria. Annu Rev Microbiol 61: 215–236. http://www.annualreviews.org/doi/10.1146/annurev.micro.61.080706.093445.

Helmann JD. 2019. Where to begin? Sigma factors and the selectivity of transcription initiation in bacteria. Mol Microbiol 112.

Hollands K, Proshkin S, Sklyarova S, Epshtein V, Mironov A, Nudler E, Groisman EA. 2012. Riboswitch control of Rho-dependent transcription termination. Proc Natl Acad Sci U S A.

Höper D, Völker U, Hecker M. 2005. Comprehensive characterization of the contribution of individual SigB-dependent general stress genes to stress resistance of Bacillus subtilis. J Bacteriol 187: 2810–2826.

Ingolia NT, Ghaemmaghami S, Newman JRS, Weissman JS. 2009. Genome-wide analysis in vivo of translation with nucleotide resolution using ribosome profiling. Science 324: 218–223. http://www.sciencemag.org/content/324/5924/218.full.pdf.

Kudla G, Murray AW, Tollervey D, Plotkin JB. 2009. Coding-sequence determinants of expression in escherichia coli. Science (80-) 324.

Lalanne J, Parker DJ, Li G. 2020. Spurious regulatory connections dictate the expression-fitness landscape of translation termination factors. bioRxiv 1–25.

Lalanne JB, Taggart JC, Guo MS, Herzel L, Schieler A, Li GW. 2018. Evolutionary Convergence of Pathway-Specific Enzyme Expression Stoichiometry. Cell 749–761.

Li GW. 2015. How do bacteria tune translation efficiency? Curr Opin Microbiol 24.

Li GW, Burkhardt D, Gross C, Weissman JS. 2014. Quantifying absolute protein synthesis rates reveals principles underlying allocation of cellular resources. Cell 157: 624–635. http://dx.doi.org/10.1016/j.cell.2014.02.033.

Lodish HF. 1968. Bacteriophage f2 RNA: Control of translation and gene order. Nature 220.

Lorenz R, Tafer H, Höner zu Siederdissen C, Stadler PF, Bernhart SH, Hofacker IL, Flamm C. 2011. ViennaRNA Package 2.0. Algorithms Mol Biol 6: 26.

Mustoe AM, Busan S, Rice GM, Hajdin CE, Peterson BK, Ruda VM, Kubica N, Nutiu R, Baryza JL, Weeks KM. 2018. Pervasive Regulatory Functions of mRNA Structure Revealed by High-Resolution SHAPE Probing. Cell 173: 181–195.e18. http://dx.doi.org/10.1016/j.cell.2018.02.034.

Nicolas P, Mäder U, Dervyn E, Rochat T, Leduc A, Pigeonneau N, Bidnenko E, Marchadier E, Hoebeke M, Aymerich S, et al. 2012. Condition-dependent transcriptome reveals high-level regulatory architecture in Bacillus subtilis. Science 335: 1103–6. http://classic.sciencemag.org/content/335/6072/1103.full.

Paget MS. 2015. Bacterial sigma factors and anti-sigma factors: Structure, function and distribution. Biomolecules 5.

Parker DJ, Lalanne J-B, Kimura S, Johnson GE, Waldor MK, Li G-W. 2020. Growth-Optimized Aminoacyl-tRNA Synthetase Levels Prevent Maximal tRNA Charging. Cell Syst 1–10.

Petersohn A, Bernhardt J, Gerth U, Höper D, Koburger T, Völker U, Hecker M. 1999. Identification of σ(B)-dependent genes in Bacillus subtilis using a promoter consensus-directed search and oligonucleotide hybridization. J Bacteriol.

Price CW. 2014. General Stress Response in Bacillus subtilis and Related Gram-Positive Bacteria. In Bacterial Stress Responses, pp. 301–318, ASM Press, Washington, DC, USA.

Reuter JS, Mathews DH. 2010. RNAstructure: Software for RNA secondary structure prediction and analysis. BMC Bioinformatics 11.

Rouskin S, Zubradt M, Washietl S, Kellis M, Weissman JS. 2014. Genome-wide probing of RNA structure reveals active unfolding of mRNA structures in vivo. Nature 505: 701–5. http://www.ncbi.nlm.nih.gov/pubmed/24336214.

Shine J, Dalgarno L. 1974. The 3’ terminal sequence of Escherichia coli 16S ribosomal RNA: complementarity to nonsense triplets and ribosome binding sites. Proc Natl Acad Sci U S A 71: 1342–1346.

Spitale RC, Flynn RA, Zhang QC, Crisalli P, Lee B, Jung JW, Kuchelmeister HY, Batista PJ, Torre EA, Kool ET, et al. 2015. Structural imprints in vivo decode RNA regulatory mechanisms. Nature 519.

Takyar S, Hickerson RP, Noller HF. 2005. mRNA helicase activity of the ribosome. Cell 120.

Tomezsko PJ, Corbin VDA, Gupta P, Swaminathan H, Glasgow M, Persad S, Edwards MD, Mcintosh L, Papenfuss AT, Emery A, et al. 2020. Determination of RNA structural diversity and its role in HIV-1 RNA splicing. Nature.

Wen J-D, Lancaster L, Hodges C, Zeri A-C, Yoshimura SH, Noller HF, Bustamante C, Tinoco I. 2008. Following translation by single ribosomes one codon at a time. Nature 452: 598–603. http://www.nature.com/nature/journal/v452/n7187/pdf/nature06716.pdf.

Yakhnin H, Pandit P, Petty TJ, Baker CS, Romeo T, Babitzke P. 2007. CsrA of Bacillus subtilis regulates translation initiation of the gene encoding the flagellin protein (hag) by blocking ribosome binding. Mol Microbiol.

Yakhnin H, Zhang H, Yakhnin A V., Babitzke P. 2004. The trp RNA-Binding Attenuation Protein of Bacillus subtilis Regulates Translation of the Tryptophan Transport Gene trpP (yhaG) by Blocking Ribosome Binding. J Bacteriol.

Zhu B, Stülke J. 2018. SubtiWiki in 2018: From genes and proteins to functional network annotation of the model organism Bacillus subtilis. Nucleic Acids Res 46: D743–D748.

Zubradt M, Gupta P, Persad S, Lambowitz AM, Weissman JS, Rouskin S. 2016. DMS-MaPseq for genome-wide or targeted RNA structure probing in vivo. Nat Methods 14: 75–82. http://dx.doi.org/10.1038/nmeth.4057.

