## Supplemental Figure 1 for "Translational activation by an alternative sigma factor in *Bacillus subtilis*"

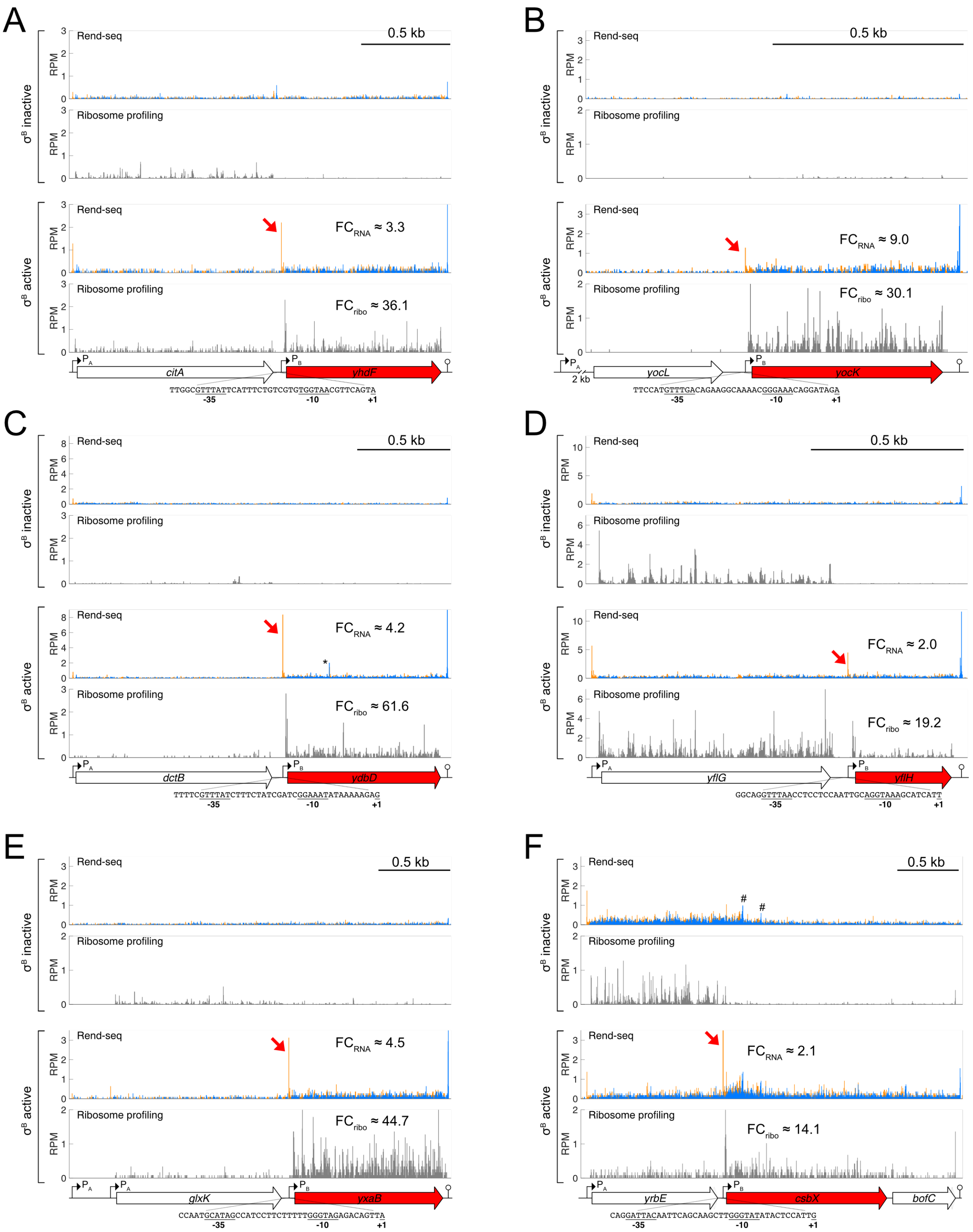
A
B
C
D
E
F
σB inactive
σB active
Rend-seq
Ribosome profiling
RPM
0.5 kb
FCRNA ≈ 3.3
FCribo ≈ 36.1
citA
yhdF
TTGGCGTTTATTCATTTCTGTCGTGTGGTAACGTTCAGTA
FCRNA ≈ 9.0
FCribo ≈ 30.1
2 kb
yocL
yocK
TTCCATGTTTGACAGAAGGCAAAACGGGAAACAGGATAGA
FCRNA ≈ 4.2
FCribo ≈ 61.6
dctB
ydbD
TTTTCGTTTATCTTTCTATCGATCGGAAATATAAAAAGAG
FCRNA ≈ 2.0
FCribo ≈ 19.2
yflG
yflH
GGCAGGTTTAACCTCCTCCAATTGCAGGTAAAGCATCATT
FCRNA ≈ 4.5
FCribo ≈ 44.7
glxK
yxaB
CCAATGCATAGCCATCCTTCTTTTTGGGTAGAGACAGTTA
FCRNA ≈ 2.1
FCribo ≈ 14.1
yrbE
csbX
bofC
CAGGATTACAATTCAGCAAGCTTGGGTATATACTCCATTG
PA
PB
-35
-10
+1

1

**Figure S1. Additional translationally activated genes with $\sigma^B$ transcript inside $\sigma^A$ polycistron.**

Rend-seq and ribosome profiling data from conditions with inactive/active $\sigma^B$ for the operons containing (**A**) *yhdF*, (**B**) *yocK*, (**C**) *ydbD*, (**D**) *yflH*, and (**E**) *yxaB*, (**F**) *csbX*. Orange and blue bars represent 5'- and 3'-mapped read counts, respectively, and the black scale bars correspond to 0.5 kb. Rend-seq data indicating the $\sigma^B$ transcription start sites are marked by red arrows. Fold changes (FC) between $\sigma^B$ active and $\sigma^B$ inactive conditions are shown. Putative $\sigma^B$-dependent promoter sequences are indicated for each gene (+1 corresponds to the 5' end of the $\sigma^B$-dependent isoform mapped by Rend-seq). The *yocK* $\sigma^A$ polycistron includes multiple upstream genes not shown for clarity. A likely post-lysis RNase A 5'/3' doublet in *ydbD* is marked by *. PnpA and Rho dependent 3' ends (Lalanne et al., 2018) in *csbX* are indicated with #. See also Figure 2.
