## Supplemental Figure 2 for "Translational activation by an alternative sigma factor in *Bacillus subtilis*"

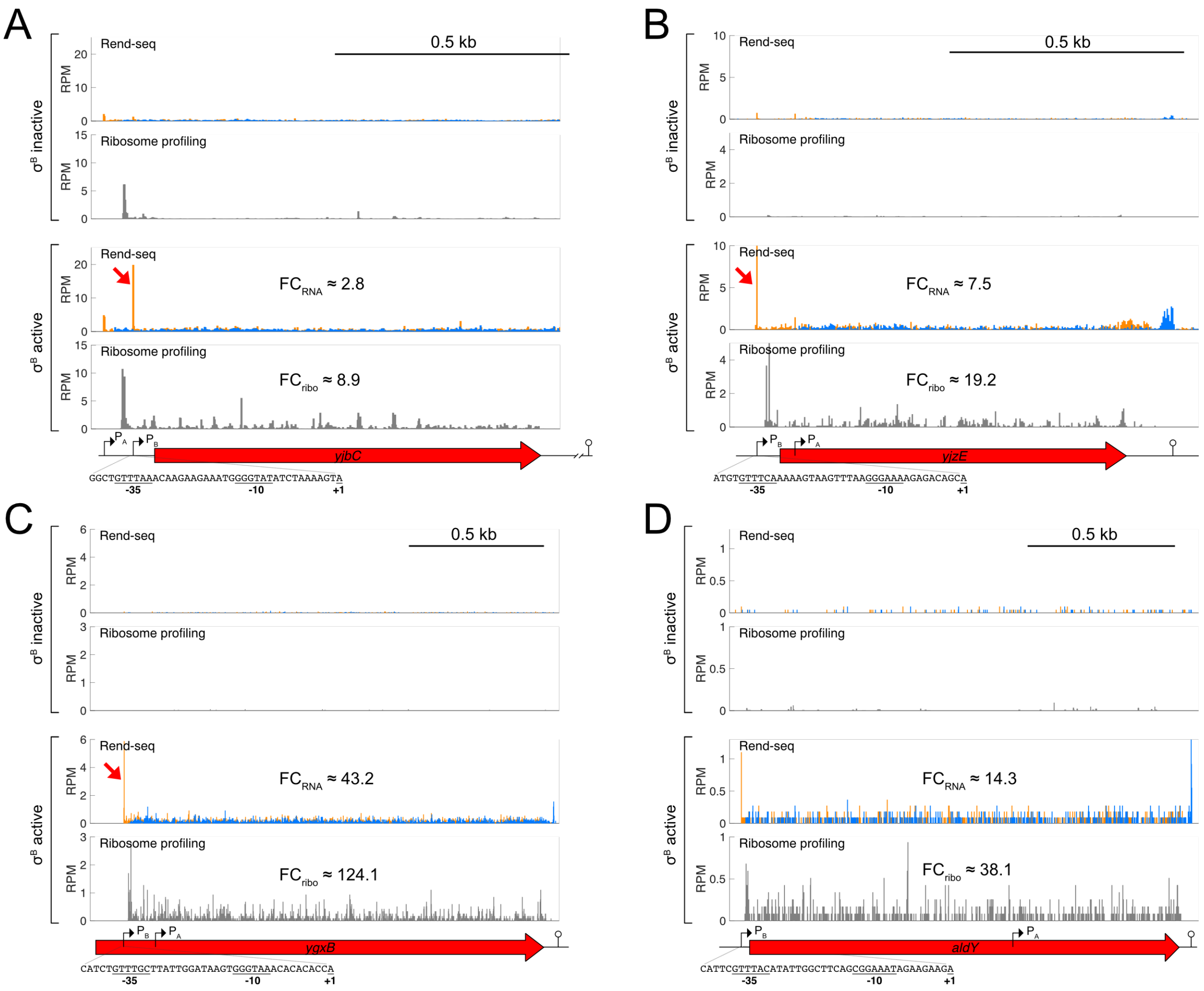


**Figure S2. Additional $\sigma^B$ translationally activated genes with changes in RNA isoform.** Rend-seq and ribosome profiling data from conditions with inactive/active $\sigma^B$ for the operons containing (**A**) *yjbC*, (**B**) *yjzE*, (**C**) *ygxB*, (**D**) *aldY*. Orange and blue bars represent 5'- and 3'-mapped read counts, respectively, and the black scale bars correspond to 0.5 kb. Rend-seq data indicating the $\sigma^B$ transcription start sites are marked by red arrows. Fold changes (FC) between $\sigma^B$ active and $\sigma^B$ inactive conditions are shown. Putative $\sigma^B$-dependent promoter sequences are indicated for each gene (+1 corresponds to the 5' end of the $\sigma^B$-dependent isoform mapped by Rend-seq). *yjbC*'s mRNA is part of a polycistronic transcript including *spxA*, which is not shown for clarity. Due to low sequencing coverage, Rend-seq data from (Lalanne et al., 2018) was used to annotate some transcript isoforms in the $\sigma^B$ inactive condition. We note that the annotation for *ygxB* is likely incorrect based on our ribosome profiling data.
