## Supplemental Figure 3 for "Translational activation by an alternative sigma factor in *Bacillus subtilis*"

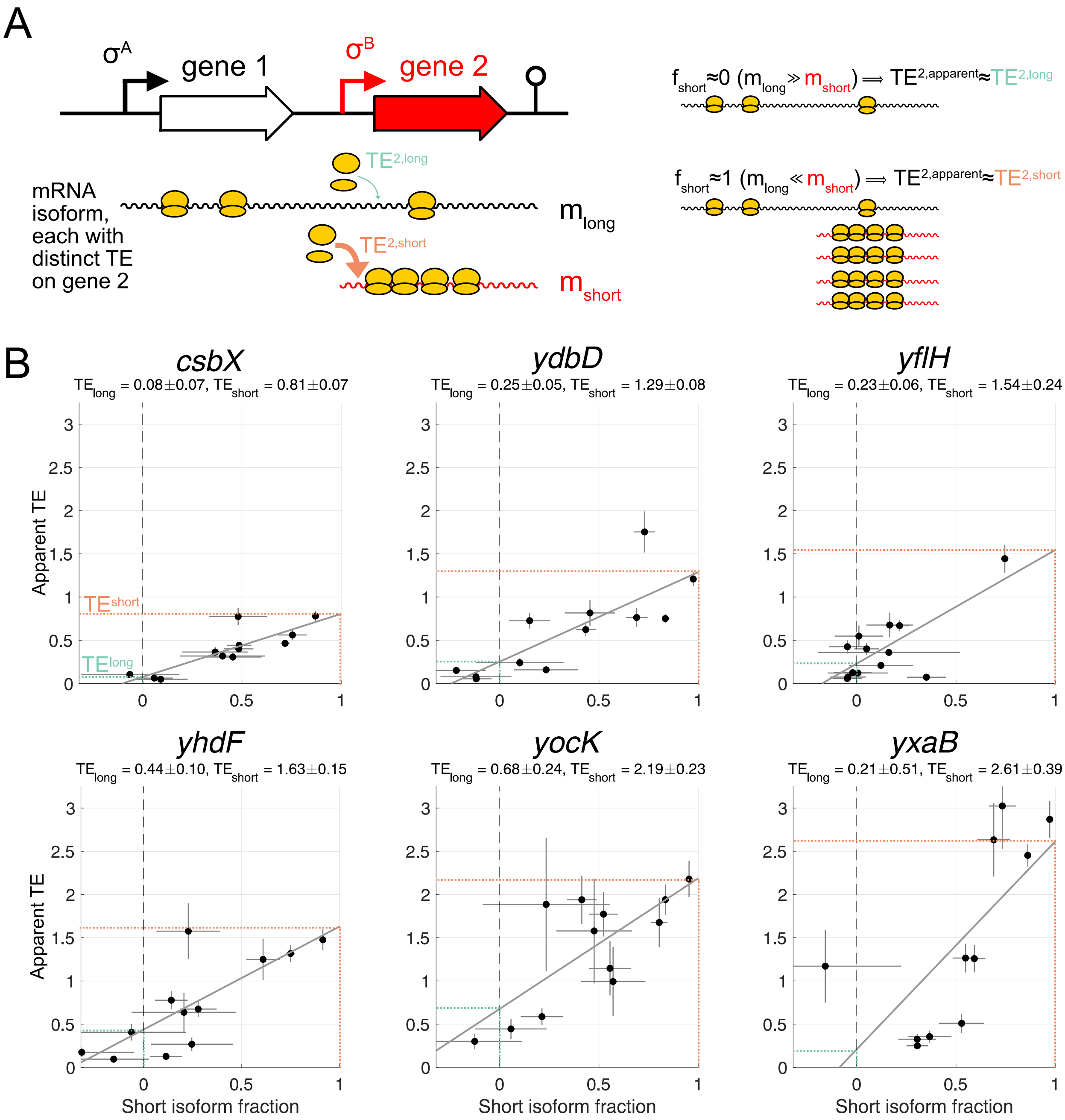
A
σA
gene 1
σB
gene 2
TE2,long
mRNA isoform, each with distinct TE on gene 2
mlong
TE2,short
mshort
fshort≈0 (mlong≫mshort) ⟹ TE2,apparent≈TE2,long
fshort≈1 (mlong≪mshort) ⟹ TE2,apparent≈TE2,short
B
csbX
TElong = 0.08±0.07, TEshort = 0.81±0.07
ydbD
TElong = 0.25±0.05, TEshort = 1.29±0.08
yflH
TElong = 0.23±0.06, TEshort = 1.54±0.24
yhdF
TElong = 0.44±0.10, TEshort = 1.63±0.15
yocK
TElong = 0.68±0.24, TEshort = 2.19±0.23
yxaB
TElong = 0.21±0.51, TEshort = 2.61±0.39
Apparent TE
TEshort
TElong
Short isoform fraction

**Figure S3. Estimations of isoform-specific TE for additional translationally activated $\sigma^B$ regulon genes.**

(**A**) Schematic illustration of the mathematical analysis used to identify the isoform-specific TE.

(**B**) Estimation of the isoform-specific TE for the short, $\sigma^A$-dependent and long, $\sigma^B$-dependent isoforms of *csbX*, *ydbD*, *yflH*, *yhdF*, *yocK*, and *yxaB*. Each point is an experimental condition which has a different short isoform fraction and correspondingly different apparent TE. Error bars correspond to standard deviations from subsampling bootstraps. The gray lines are linear regressions, whereas the dashed lines indicate estimates of isoform-specific TE calculated from the fits (Methods). Estimated isoform-specific TEs and errors (standard deviations) from a bootstrapped linear fit (Methods) are shown. See also Figure 3.
